# Insights into ontogenetic scaling and morphological variation in sharks from near-term brown smooth-hound (*Mustelus henlei*) embryos

**DOI:** 10.1101/2024.02.05.578906

**Authors:** Joel H. Gayford, Phillip C. Sternes, Scott G. Seamone, Hana Godfrey, Darren A. Whitehead

## Abstract

Elasmobranchs (sharks and rays) exhibit a wide range of body forms adapted to various ecological niches. Body form differs not only between species, but between life stages of individual species as a result of ontogenetic allometry. In sharks, it has been proposed that these ontogenetic shifts in body form result from shifts in trophic and/or spatial ecology (the allometric niche shift hypothesis). Alternatively, it has been suggested that ontogenetic allometry may result from intrinsic morphological constraints associated with increasing body size – e.g. to counteract shifts in form-function relationships that occur as a function of size and could compromise locomotory performance. One major limitation affecting our understanding of ontogenetic scaling in sharks is that existing studies focus on postpartum ontogeny – ignoring the period of growth that occurs prior to birth/hatching. In this study, we report ontogenetic growth trajectories from 39 near-term brown smooth hound (*Mustelus henlei*) embryos taken from manually collected measurements. We found that unlike most other species and later ontogenetic stages of *M. henlei*, these embryos predominantly grow isometrically, and appear to display relatively high levels of morphological disparity. These results provide rudimentary support for the allometric niche shift hypothesis (as in the absence of ontogenetic niche shifts isometry dominates body-form scaling) and provide important insight into early shark ontogeny and morphological/developmental evolution.

## Introduction

Elasmobranchs (sharks and rays) are a large radiation of marine vertebrates that have persisted through periods of intense environmental change and represent an ecologically important component of marine diversity (Heithaus et al., 2010; Grogan et al., 2012; Flowers et al., 2021). Elasmobranchs are extremely diverse in terms of both morphology and ecology (Maisey et al., 2004; Navia et al., 2007; Cortés et al., 2008; Sternes and Shimada, 2020; Kuraku, 2021; Mull et al., 2022; Andrzejaczek et al., 2022). Whilst many knowledge gaps remain regarding our understanding of elasmobranch ecology, it has long been known that some species exhibit shifts in spatial and trophic ecology through development (Grubbs, 2010). It is only relatively recently however that ontogenetic shifts in morphology have been documented (e.g. Lingham-Soliar, 2005; Irschick and Hammerschlag, 2015; Fu et al., 2016; Irschick et al., 2017; Ahnelt et al., 2020; Sternes and Higham, 2022; Bellodi et al., 2023; Gayford et al., 2023a; Gayford et al., 2023b; Seamone et al., 2023; Yun and Watanabe, 2023; Gayford et al., 2024). These shifts are important not only from the perspective of functional ecology but help us to understand the selective forces underlying the evolution of elasmobranch morphology (Gayford et al., 2023b). Various hypotheses exist for observed growth trajectories, including selection relating to spatial and trophic ecology and fundamental constraints on locomotor performance associated with increasing body size (Irschick et al., 2017; Gayford et al., 2023b; Seamone et al., 2023). Typically, a mosaic of allometric and isometric growth (where some structure grows disproportionately or proportionately relative to body size respectively) is observed in functionally important morphological measurements, with the nature of these scaling relationships varying between size classes and sexes in some cases (Gayford et al., 2023b). Despite the increasing number of studies addressing the topic of ontogenetic allometry in a range of shark species, there exist a number of factors that severely limit our understanding of the phenomenon, including taxonomic coverage, sample sizes, and an understanding of the genetic/developmental and functional basis of the morphological structures in question (Sternes and Higham, 2022; Gayford, 2023; Gayford et al., 2023a).

One potential limitation that has not yet been addressed in the literature is that existing ecomorphological studies of scaling in elasmobranchs focus predominantly on postnatal ontogeny, despite the evolutionary and ecological significance of morphological changes occurring during prenatal ontogeny. Those studies that do address embryogenic scaling focus on developing staging tables without much focus on the ecomorphology of scaling itself (Tomita et al., 2018; López-Romero et al., 2020; Byrum et al., 2023). They may, as in other ontogenetic stages (Gayford et al., 2023b) act to maximise fitness in the context of the trophic and spatial ecology of this taxon post-partum. It is also important to consider that embryo morphology may also be influenced by prepartum environmental conditions (Kaplan and Phillips, 2006; Rodda and Seymour, 2008). Selective pressures associated with prepartum conditions could include sibling relatedness (Pfennig and Collins, 1993), constraints relating to the anatomy of the mother (Qualls et al., 1995), and even temporal variation in the environmental conditions experienced by the mother (Sale et al., 2007; McCoy et al., 2020). Each of these factors is yet to be considered from the perspective of elasmobranch taxa, representing a significant gap in our understanding of morphological evolution within this clade. Studies of scaling in embryos may also provide insight into the selective pressures driving the evolution of allometric and isometric growth, as shark embryos typically exhibit substantial proportional body-size changes (Tomita et al., 2018; López-Romero et al., 2020; Byrum et al., 2023) without undergoing major shifts in the trophic/spatial environment.

The prepartum environment is important to the process of development across taxa, but is arguably of particular interest in elasmobranchs due to the complexity of their reproductive biology: not only are multiple mating systems observed amongst extant elasmobranch taxa (Bester-van der Merwe et al., 2022), but substantial variation has been reported in parameters such as brood size, gestation periods, ovarian cycles and reproductive behaviours (Carrier et al., 2004). Elasmobranch reproduction is a complex arena of intense genetic conflict between multiple players: sexually antagonistic coevolution (Portnoy and Heist, 2012), male-male conflict (Rowley et al., 2019) and sibling conflict (Chapman et al., 2013) have all been shown to occur within this clade and have potential consequences for morphological evolution. Sexually antagonistic coevolution has been attributed with the evolution of a number of sexual dimorphisms (Kajiura and Tricas, 1996; Whitehead et al., 2022), and intrauterine cannibalism (one manifestation of sibling conflict) (Gilmore et al., 2011) would logically impart strong selection pressures on tooth morphology and the developmental timing of tooth acquisition.

The brown smoothhound shark, *Mustelus henlei* (Gill, 1863) is a primarily demersal Carcharhiniform shark found along the Pacific coast of the Americas, from California to Peru (Compagno, 1984; Ebert et al., 2021). This species is heavily fished in Baja California, Mexico (Medina-Morales et al., 2020; Smith et al., 2009), which has facilitated several studies into its biology and ecology in the region (Pérez-Jiménez and Sosa-Nishizaki, 2008; Byrne and Avise, 2012; Pantoja-Echevarría et al., 2020; Gayford et al., 2023a). *M. henlei* has a relatively great distribution compared to other *Mustelus* species with which it exhibits some degree of range overlap (Chabot et al., 2015), however it appears that populations are differentiated both genetically (Chabot and Haggin, 2014; Chabot et al., 2015) and ecologically, with ontogenetic and sex-based trophic variation, and overall trophic position appearing to differ between study sites (Ruso, 1975; Espinoza et al., 2012; Rodríguez-Romero et al., 2013; Amariles et al., 2017). There is a paucity of data regarding reproductive biology in this species (Pérez-Jiménez and Sosa-Nishizaki, 2008). Multiple paternity is known to be present in *M. henlei* (Chabot and Haggin, 2014; Rendón-Herrera et al., 2022), and basic information exists regarding observations of placental viviparity and variation in litter size (Yudin, 1987; Pérez-Jiménez and Sosa-Nishizaki, 2008), however no data exist regarding the morphogenesis or early ontogeny of *M. henlei*. A recent study did present ontogenetic growth trajectories for this species, using a large dataset of both adult and juvenile (but not neonate or embryonic) specimens (Gayford et al., 2023a). For this reason, *M. henlei* presents an ideal case study through which to compare ontogenetic growth trajectories in pre and postpartum environments.

In this study, we use measurements obtained from *M. henlei* embryos to explore possible ontogenetic shifts in body form during this crucial stage of development. We seek to compare growth trajectories for various morphological structures with those found from post-natal individuals (both juvenile and adult) and propose potential explanations for any differences observed. This study provides a vital contribution to existing literature on ontogenetic morphological shifts in elasmobranch taxa as existing evolutionary studies have focussed on post-natal ontogeny – neglecting the potential selective influence of the pre-natal environment on morphology and growth trajectories.

## Methodology

### Ethics statement

Data collection and analysis procedures in this study complied with Mexican animal welfare laws and guidelines. No permit or ethical approval was necessary as animals were caught as part of legal artisanal fisheries and data were only collected after landing with permission of the fishers. Participants of this study neither promote nor encourage the harvesting of sharks.

### Data Collection

*M. henlei* comprise a major component of total catch at artisanal fishing camps on the Pacific coast of Baja California Sur, Mexico. During the month of December 2022, we recovered paired uteri from 5 pregnant females at an artisanal fish camp. Full uteri were recovered (Figure 1), however due to the rate at which sharks are processed at this camp, it was not possible to match uteri to specific adult females. Uteri were stored at 5°C and transported to a laboratory for analysis within 24 hours. As we retained full uteri, embryos did not desiccate and therefore did not exhibit any significant morphological deformation. At the laboratory, uteri were dissected and embryos where all morphological structures of interest were clearly present were isolated for data collection. Damaged embryos, or those too small for morphological structures to be reliably measured were not included. From each of the remaining 39 embryos, 28 morphological measurements capturing variation in body form were taken recorded with a tape measure to the nearest millimetre (Table 1).

**Table 1:** Morphological measurements taken from embryos, their abbreviations and definitions.

| Character | Abbreviation | Definition |
| --- | --- | --- |
| Total length | TL | Distance from the tip of the snout to the tip of the upper caudal lobe |
| Precaudal length | PL | Distance from the tip of the snout to the precaudal pit |
| Upper caudal length | UL | Distance from the upper insertion point of the caudal fin to the tip of the caudal fin upper lobe (posterior tip) |
| Lower caudal length | LL | Distance from the lower insertion point of the caudal fin to the tip of the caudal fin lower lobe (ventral tip) |
| Caudal height | CH | Distance between the posterior tip and ventral tip of the caudal fin |
| Caudal keel | CK | Total circumference at the caudal keel |
| Lateral span | LS | Distance over the dorsal body surface, measured between the anterior insertion points of the left and right pectoral fins |
| Frontal span | FS | Distance over the dorsal body surface at the anterior edge of the insertion point of the first dorsal fin, measured between points on the flank on the same horizontal plane as the pectoral fin insertion points |
| Proximal span | PS | Distance over the dorsal body surface at the posterior edge of the insertion point of the first dorsal fin, measured between points on the flank on the same horizontal plane as the pectoral fin insertion points |
| Frontal span 2 | FS2 | Distance over the dorsal body surface at the anterior edge of the insertion point of the second dorsal fin, measured between points on the flank on the same horizontal plane as the pelvic fin insertion points |
| Proximal Span 2 | PS2 | Distance over the dorsal body surface at the posterior edge of the insertion point of the second dorsal fin, measured between points on the flank on the same horizontal plane as the pelvic fin insertion points |
| Dorsal length | DL | Distance from the anterior insertion point of the first dorsal fin to the upper tip of the same fin |
| Dorsal width | DW | Horizontal distance from the anterior insertion point of the first dorsal fin to the posterior tip of the same fin |
| Dorsal height | DH | Vertical distance from the tip of the first dorsal fin to the base of the first dorsal fin |
| Second dorsal length | DL2 | Distance from the anterior insertion point of the second dorsal fin to the upper tip of the same fin |
| Second dorsal width | DW2 | Horizontal distance from the anterior insertion point of the first dorsal fin to the posterior tip of the same fin |
| Second dorsal height | DH2 | Vertical distance from the tip of the first dorsal fin to the base of the second dorsal fin |
| Pectoral fin length | PF | Distance from the distal insertion point of the pectoral fin to the fully extended tip of the pectoral fin |
| Pectoral fin width | PFW | Distance from the proximal insertion point of the pectoral fin to the distal insertion point of the pectoral fin |
| Pelvic fin length | PLF | Distance from the distal insertion point of the pelvic fin to the fully extended tip of the pelvic fin |
| Pelvic fin width | PLFW | Distance from the proximal insertion point of the pelvic fin to the distal insertion point of the pelvic fin |
| Anal fin length | AL | Distance from the anterior insertion point of the anal fin to the upper tip of the same fin |
| Anal fin height | AH | Horizontal distance from the anterior insertion point of the anal fin to the posterior tip of the same fin |
| Anal fin width | AW | Vertical distance from the tip of the anal fin to the base of the first dorsal fin |
| Eye to eye | EE | Distance over the dorsal body surface from the midpoint of one eye to the midpoint of the other |
| Eye height | EH | Vertical diameter of the eye at its midpoint |
| Eye length | EL | Horizontal diameter of the eye at its midpoint |
| Mouth diameter | MD | Linear distance over the ventral body surface between the posterior most insertion points of the lower jaw |

**Figure 1:**
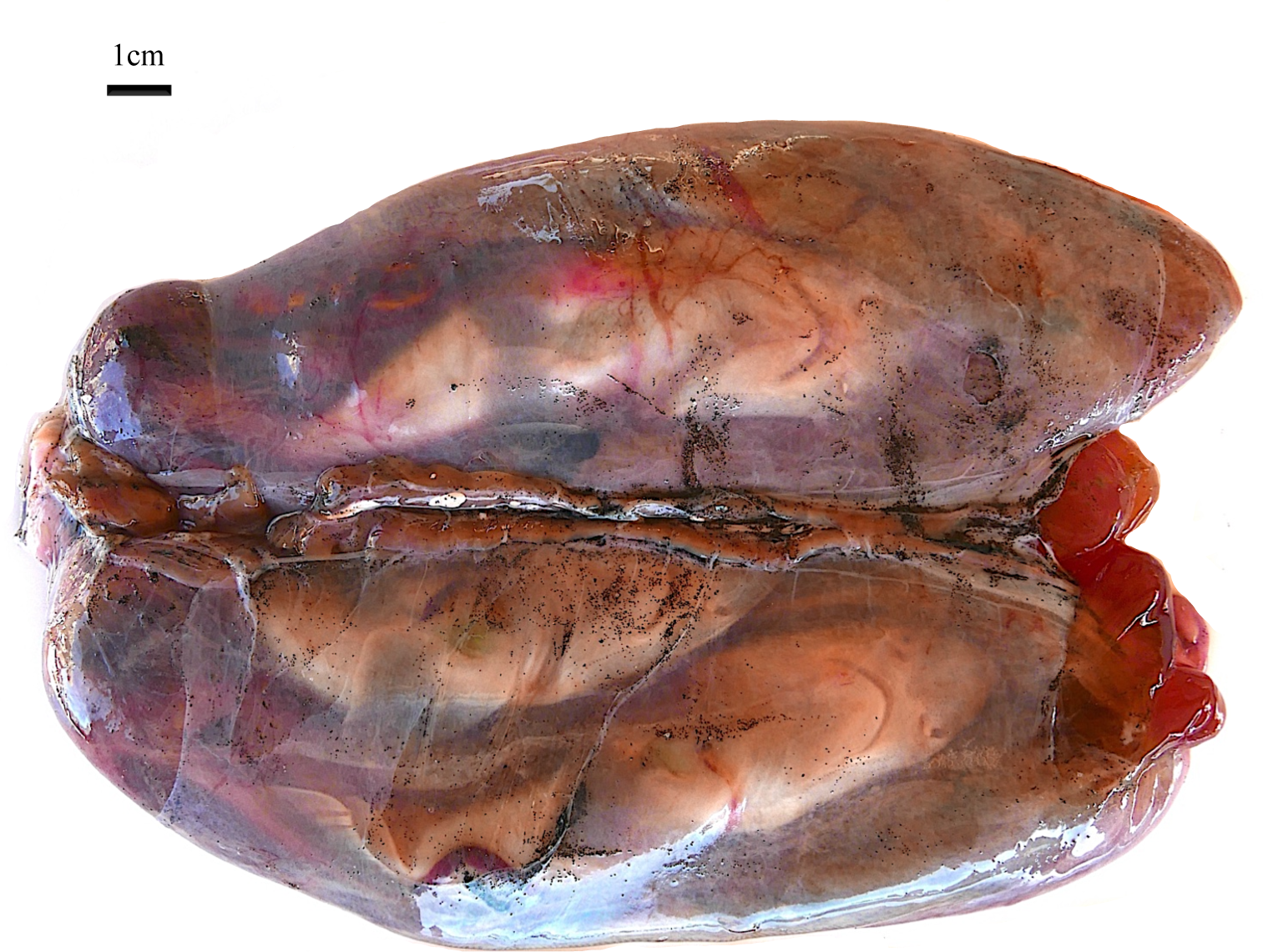
Paired uteri from one pregnant M. henlei individual, prior to extraction and measurement of embryos.

### Data analysis

No developmental staging table exists for *M. henlei*. To assess the general developmental stage of embryos, external morphology was compared to that described in embryological scaling tables for other shark species (e.g. Ballard et al., 1993; Rodda and Seymour, 2008; Onimaru et al., 2018).

To extract ontogenetic growth trajectories for morphological measurements, simple linear regression analysis was performed using the function *lm* in the R statistical environment (R Core Team, 2023). All data were log-transformed prior to statistical analyses, such that growth curves are represented by linear (rather than exponential) relationships, the gradient of which determines whether growth is allometric or isometric. Each measurement (excluding TL and PL) was regressed separately against precaudal length (PL). The observed scaling coefficient (gradient) was then compared to a null hypothesis of isometry (equivalent to a gradient of 1.00). Where scaling coefficients are found to be significantly (p<0.05) lower or higher than 1.00 they represent negative and positive allometric relationships respectively (the absence of any significant relationship is hence an extreme case of negative allometry). This means that as body size increases, the measurement in question becomes disproportionately smaller or larger, respectively.

## Results

Embryos ranges in size from 168mm to 208mm in total length (Table 2), within the size range at which pups are born (Ebert et al., 2021). All embryos were recovered in good condition, enabling all measurements to be taken from all individuals. The largest embryos were determined to be near term (close to parturition) on the basis of their well-developed external morphology, and comparison with existing developmental staging tables for other shark species (Ballard et al., 1993; Rodda and Seymour, 2008; Onimaru et al., 2018). Due to the lack of any published staging table for any member of the *Mustelus* genus we were unable to ascribe specific stages to each embryo.

**Table 2:** Summary statistics for all measurements included in the study, recorded to the nearest mm.

| Character | Minimum<br>(mm) | Maximum<br>(mm) | Mean<br>(mm) | Standard<br>Deviation<br>(mm) |
| --- | --- | --- | --- | --- |
| TL | 168 | 208 | 188 | 10.1 |
| PL | 138 | 168 | 151 | 7.8 |
| UL | 30 | 41 | 37 | 3.0 |
| LL | 11 | 18 | 15 | 1.6 |
| CH | 25 | 39 | 31 | 3.2 |
| CK | 7 | 19 | 13 | 2.5 |
| LS | 26 | 36 | 30 | 2.5 |
| FS | 19 | 29 | 23 | 2.6 |
| PS | 11 | 22 | 18 | 2.0 |
| FS2 | 8 | 16 | 13 | 1.6 |
| PS2 | 7 | 12 | 10 | 1.2 |
| DL | 17 | 24 | 21 | 1.7 |
| DW | 20 | 28 | 24 | 2.2 |
| DH | 10 | 18 | 14 | 2.1 |
| DL2 | 12 | 19 | 16 | 1.9 |
| DW2 | 14 | 23 | 19 | 1.9 |
| DH2 | 5 | 14 | 10 | 1.7 |
| PF | 19 | 26 | 22 | 1.7 |
| PFW | 13 | 22 | 18 | 2.0 |
| PLF | 10 | 14 | 12 | 1.3 |
| PLFW | 8 | 13 | 11 | 1.2 |
| AL | 8 | 12 | 10 | 1.0 |
| AH | 3 | 6 | 5 | 0.8 |
| AW | 11 | 16 | 13 | 1.2 |
| EE | 13 | 20 | 17 | 1.7 |
| EH | 3 | 4 | 3 | 0.5 |
| EL | 6 | 9 | 8 | 0.8 |
| MD | 11 | 15 | 13 | 1.2 |

Linear regression of 26 morphological measurements against precaudal length (PL) failed to identify statistically significant departures from isometry in most cases: only three measurements demonstrated allometric trajectories (PFW, AL and EL) whereas the remaining 23 measurements scale isometrically (Table 3). Each of the three allometric relationships were negative, meaning that they become disproportionately smaller through ontogeny (Table 3). Whilst the slope of the scaling relationship in these cases is significantly lower than 1, this does not rule out the possibility that there is no significant relationship between body size at PFW, AL, or EL. Indeed the low ranges for AL and EL (Table 2) relative to the measurement resolution (1mm) mean that scaling relationships for these measurements should be interpreted with caution. The proportion of variance explained by scaling relationships was low in many cases, ranging between 0.03 and 0.35 (Table 3).

**Table 3:** Linear regression results for the Log10 transformed total dataset (n=39). Significant (allometric) results (p≤0.05) are indicated in bold.

| Character | Coefficient | Std. error | t value | p value | Residual std. error | $R^2$ | Adj. $R^2$ | F value |
| --- | --- | --- | --- | --- | --- | --- | --- | --- |
| UL | 0.9565 | 0.2124 | 0.205 | 0.839 | 0.02965 | 0.354 | 0.3366 | 20.28 |
| LL | 0.9824 | 0.3135 | 0.056 | 0.956 | 0.04377 | 0.2097 | 0.1884 | 9.819 |
| CH | 1.0991 | 0.2592 | 0.382 | 0.704 | 0.03618 | 0.3271 | 0.3089 | 17.98 |
| CK | 1.3668 | 0.6503 | 0.564 | 0.576 | 0.09078 | 0.1067 | 0.0825 | 4.418 |
| LS | 0.9182 | 0.2155 | -0.380 | 0.706 | 0.03008 | 0.3292 | 0.3111 | 18.16 |
| FS | 0.9785 | 0.3003 | 0.072 | 0.943 | 0.04192 | 0.223 | 0.202 | 10.62 |
| PS | 1.0135 | 0.3426 | 0.040 | 0.969 | 0.04783 | 0.1913 | 0.1694 | 8.75 |
| FS2 | 0.7742 | 0.3945 | 0.572 | 0.571 | 0.05507 | 0.0943 | 0.0698 | 3.851 |
| PS2 | 0.4952 | 0.4184 | 1.206 | 0.235 | 0.0584 | 0.03649 | 0.0104 | 1.401 |
| DL | 0.6903 | 0.2330 | 1.329 | 0.192 | 0.03253 | 0.1917 | 0.1699 | 8.777 |
| DW | 0.8366 | 0.2525 | 0.647 | 0.522 | 0.03524 | 0.2288 | 0.208 | 10.98 |
| DH | 1.1942 | 0.4280 | 0.454 | 0.653 | 0.05975 | 0.1738 | 0.1515 | 7.785 |
| DL2 | 0.6289 | 0.3686 | 1.007 | 0.321 | 0.05146 | 0.07293 | 0.0479 | 2.91 |
| DW2 | 0.9284 | 0.2888 | 0.248 | 0.806 | 0.04032 | 0.2183 | 0.1972 | 10.33 |
| DH2 | 0.5780 | 0.5947 | 0.710 | 0.482 | 0.08302 | 0.02489 | -0.002 | 0.944 |
| PF | 0.59619 | 0.21537 | 1.875 | 0.0687 | 0.03006 | 0.1716 | 0.1492 | 7.663 |
| **PFW** | **0.1363** | **0.3663** | **2.358** | **0.0238** | **0.05113** | **0.00373** | **-0.023** | **0.139** |
| PLF | 0.4274 | 0.3416 | 1.676 | 0.102 | 0.04769 | 0.04058 | 0.0147 | 1.565 |
| PLFW | 0.5790 | 0.3545 | 1.187 | 0.243 | 0.04949 | 0.06724 | 0.0420 | 2.667 |
| **AL** | **0.3056** | **0.3104** | **2.237** | **0.0314** | **0.04334** | **0.02553** | **-0.001** | **0.969** |
| AH | 0.6326 | 0.5550 | 0.662 | 0.512 | 0.07747 | 0.03392 | 0.0078 | 1.299 |
| AW | 0.9035 | 0.2491 | 0.388 | 0.701 | 0.03477 | 0.2623 | 0.2423 | 13.15 |
| EE | 1.1957 | 0.2439 | 0.802 | 0.4276 | 0.03405 | 0.3938 | 0.3774 | 24.03 |
| EH | 0.1336 | 0.4465 | 1.941 | 0.060 | 0.06233 | 0.00241 | -0.025 | 0.090 |
| **EL** | **0.1579** | **0.3257** | **2.586** | **0.0138** | **0.04546** | **0.00631** | **-0.021** | **0.235** |
| MD | 0.48595 | 0.29460 | 1.745 | 0.0893 | 0.04112 | 0.0685 | 0.0433 | 2.721 |

## Discussion

A number of studies have investigated postnatal ontogenetic scaling trends in elasmobranch fishes. Whilst in some cases, specific morphological structures exhibit predominantly isometric or allometric growth respectively (Reiss and Bonnan, 2010; Irschick and Hammerschlag, 2015; Ahnelt et al., 2020), it appears that in most species body form develops through a combination of isometry and allometry (Irschick and Hammerschlag, 2015; Irschick et al., 2017; Sternes and Higham, 2022; Bellodi et al., 2023; Gayford et al., 2023a; Gayford et al., 2023b; Seamone et al., 2023). In this study we instead focus on the latter stages of prenatal ontogeny, finding that isometric growth is present across nearly all functionally important aspects of external morphology (Table 3). Moreover, even where allometry is present, *R*^2^ values are far lower than found in other studies (Table 3) – even across similar size ranges in the same species (Gayford et al., 2023a). This suggests that across a critical stage of ontogeny, the body form of *M. henlei* remains broadly unchanged and morphological variation is relatively great compared to postnatal ontogenetic stages. Here, we discuss the implications of these results for our understanding of body form evolution and ontogenetic scaling in sharks and the life history of *M. henlei*.

The prevalence of isometry as opposed to allometry in *M. henlei* embryos lends further credence to the idea that allometric shifts in shark body form result at least in part from ontogenetic niche shifts. Studies investigating ontogenetic shifts in body form in sharks typically relate these changes to differences in trophic or spatial ecology (Lingham-Soliar, 2005; Irschick and Hammerschlag, 2015; Fu et al., 2016; Irschick et al., 2017; Sternes and Higham, 2022; Gayford et al., 2023a; Gayford et al., 2023b; Seamone et al., 2023; Yun and Watanabe, 2023; Gayford et al., 2024). In many species, adults consume larger prey and spend a greater proportion of time in offshore, open-ocean environments – likely resulting in a shift in the selective pressures acting on individuals through ontogeny (Sternes and Higham, 2022; Gayford et al., 2023b). Niche shifts are not the only hypothesised drivers of allometry in sharks – there have also been suggestions that allometric growth may be more prevalent in larger-bodied species as a result of size-related physiological/locomotory constraints (Irschick et al., 2017; Seamone et al., 2023). This hypothesis suggests that allometric growth might act to maintain locomotor performance as body size increases, as opposed to modifying performance to suit different ecological conditions (Seamone et al., 2023). Our results support the allometric niche shift hypothesis given that, in the absence of an ontogenetic niche most measurement scale with isometry (Table 3). Additionally, the few cases of allometry are weak and explain a negligible proportion of variance in the data (Table 3). This is in stark contrast to later ontogenetic stages of *M. henlei*, (that are thought to undergo some degree of ontogenetic niche shift) where a much greater proportion of measurements scale with allometry, and allometric relationships explain substantial proportions of morphological variance (Gayford et al., 2023a).

The observed ∼25% increase in body size is unlikely to have similar physiological/locomotory consequences in embryos because they are not subject to the physical constraints of locomotion in water – and therefore we are unable to directly test the hypothesis that constraint associated with locomotor performance drives the evolution of allometry. However, our results (combined with those of Gayford et al., 2023a) suggest that in *M. henlei*, allometric growth is not an intrinsic consequence of increased body-size alone and support the role of ontogenetic niche shifts in selecting for the evolution of allometry.

The broad absence of allometric growth in *M. henlei* embryos does not match findings from other ontogenetic stages of the species, and could be indicative of morphological constraint imposed by the maternal environment. Embryo morphology in other taxa can be influenced by selection pressures that are unique to the prenatal environment (Kaplan and Phillips, 2006; Rodda and Seymour, 2008). There are obvious size and shape constraints relating to the internal anatomy and body size of the mother (or the eggcase in the case of oviparous taxa) that could hypothetically influence morphology. Moreover, selection relating to locomotion, predation, and nutrient acquisition clearly differs substantially between the prenatal and postnatal environments. In several shark species (including both matrotrophic and oviparous taxa), shifts in embryo morphology have been hypothesised to result from differences between these environments: in white sharks (*Carcharodon carcharias*), embryos undergo a dramatic transition from heterocercal to lunate caudal fins, with the latter is thought to be advantageous to fast-moving active predators such as young white sharks (Tomita et al., 2018). Interestingly, this transition continues well into postnatal ontogeny (Lingham-Soliar, 2005). The early development and subsequent loss of external gill filaments recorded in various elasmobranch taxa is thought to result from potential hypoxia during early stages of development, which is of course unique to the prenatal environment (Rodda and Seymour, 2008). Such shifts are clearly not present in *M. henlei*, as body form remains broadly constant through the latter stages of prenatal ontogeny (Table 3).

There are two potential explanations for why *M. henlei* predominantly exhibits isometric growth during late prenatal ontogeny: either selective constraint on embryo morphology imposed by the maternal environment prevents morphology optimised for the postnatal environment from arising until after parturition, or selective constraint imposed by the maternal environment is largely absent, and isometry results from acquisition of optimal postnatal morphology during early prenatal ontogeny. Measurements included in this study such as DH and PF define the maximum body diameter of embryos, and thus may be under constraint as they could influence the number of embryos able to fit within the uteri, or the probability of injury to the mother during parturition. In line with this, body girth measurements along the trunk grow isometrically in embryos (Table 3), despite showing significant positive allometry in later postnatal life stages (Gayford et al., 2023a). It is not however possible to conclusively distinguish between this hypothetical constraint and an ontogenetic trajectory in which the morphology favoured by selection in the postnatal environment is acquired early in prenatal ontogeny on the basis of our data. Ontogenetic morphological trajectories for neonate *M. henlei* would enable us to discern between these hypotheses, as a predominance of isometric growth here would rule out postnatal allometry as a mechanism for overcoming prenatal morphological constraint. Contrastingly, predominance of allometric growth would suggest that there are substantial differences between the body forms favoured in prenatal and postnatal environments, providing rudimentary evidence for morphological constraint.

Various theoretical models have been produced to describe patterns of morphological variation through vertebrate embryogenesis and ontogeny (Von Baer, 1828; Keibel and Abraham, 1900). Whilst not universally accepted, the ‘hourglass model’ is amongst the most supported of these models, and suggests that morphological diversity is greatest during the earliest and latest stages of embryogenesis (Irie and Kurutani, 2014; Irie, 2017). This is thought to be due to more stringent developmental constraints acting on intermediate embryogenic stages (Piasecka et al., 2013). These models have traditionally been used to compare morphological divergence between taxa, however it has previously been suggested that patterns of intraspecific morphological variation could follow an hourglass model (Pantalacci and Sémon, 2015). We cannot empirically test the validity of an hourglass model with the data in this study, nevertheless our results are consistent with the concept of relatively high morphological variance during the latter stages of embryogenesis, as evidenced by the low *R*^2^ values relative to similar proportional body size ranges in latter ontogenetic stages of the same species (Table 3; Gayford et al., 2023a). Comparing our results to these latter ontogenetic stages, this suggests that selection/constraint on morphology may be weakened or relaxed during the latter stages of embryogenesis of *M. henlei* relative to subsequent life stages (Gayford et al., 2023a). This also provides further indirect support for the niche shift-driven allometry, suggesting that where niche shifts are not present, selection on body form is relatively weak. Importantly, the evolutionary constraint posited by developmental models is not the same as the physical constraints on morphology that could theoretically influence embryo morphology mentioned previously.

Without additional genetic and morphological information from the earlier stages of embryogenesis it is impossible to prove whether variation in the intensity of developmental constraint is responsible for the morphological variation reported here. There are indeed several plausible alternative explanations for such variation. In the case of some measurements (particularly AL and EL), the measurement resolution (1mm) is fairly low relative to the measurement range (Table 2), and this could lead to spuriously low *R*^2^ values. However this is not the case for all measurements considered here (Table 2) and thus, that measurement resolution alone cannot explain this apparent high morphological variance. Multiple paternity (where a single brood consists of individuals sired by more than one male) is a known phenomenon in *M. henlei* (Byrne and Avise, 2012; Chabot and Haggin, 2014; Réndon-Herrera et al., 2022) and could hypothetically contribute to morphological variation. However, the potential influence of multiple paternity is difficult to quantify due to uncertainty regarding the genetic basis of morphological traits in sharks (Gayford, 2023), and geographic variation in the extent of multiple paternity (Chabot and Haggin, 2014; Réndon-Herrera et al., 2022). Even if the genotypes of potential sires do not result in substantial morphological variation, multiple paternity could result in morphological differences between embryos due to latency of fertilisation within a brood. In several shark species where multiple paternity is present, fertilisation is thought to occur over an extended period of time (Schmidt et al., 2010; Marino et al., 2015). In this scenario there may be discrepancies in the maternal provisioning that different embryos receive at a given embryogenic stage, as in matrotrophic species the nutrients provided to embryos by the mother depends to some extent on the trophic characteristics of the environmental conditions to which the mother is exposed (Olin et al., 2011; McCoy et al., 2020). The degree of body size variation in our results (Table 2) combined with the ∼10 month gestation period and seasonal migratory behaviour of *M. henlei* (Pérez-Jiménez and Sosa-Nishizaki, 2008) leads us to suggest that multiple paternity in this taxon likely does result in different embryos within a litter receiving different nutritional profiles from the mother at given embryogenic stages. Whether this contributes to morphological variation remains unknown, but we suggest that future studies should investigate the extent to which models of developmental constraint, multiple paternity, and delayed fertilisation could contribute to intraspecific and interspecific patterns of morphological variation.

## Conclusion

In this study we have shown that during the latter stages of prenatal ontogeny, *M. henlei* body form grows predominantly isometrically, in stark contrast with postnatal ontogenetic stages of this species. These results not only improve our understanding of the life history of *M. henlei* but provide valuable insight into the evolution of ontogenetic scaling and morphological diversity in sharks. In the context of the prenatal environment, isometric growth supports the allometric niche shift hypothesis and relatively high morphological disparity between embryos raises questions about shark reproductive biology, evolutionary genetics and development that warrant further study. Our dataset is undoubtedly limited – both in terms of ontogenetic coverage and measurement resolution. However, similar studies covering similar proportional ranges (body size increases of ∼25-50%) of body size (Irschick and Hammerschlag, 2015; Gayford et al., 2023a) have been key in forming current opinion on ontogenetic scaling in sharks. Future studies considering different ontogenetic stages of *M. henlei* and similar ontogenetic stages in other shark species will enable us to further discern between some of the hypotheses addressed in this study – particularly in the cases of developmental constraints and morphological disparity between embryos. Ultimately this study acts as a baseline against which ontogenetic body form trajectories from embryos of other shark species can be compared, within the same quantitative and theoretical framework that has been applied to studies of ontogenetic scaling in postnatal life-stages of elasmobranch taxa.

